# Limited O-specific polysaccharide (OSP)-specific functional antibody responses in young children with Shigella infection in Bangladesh

**DOI:** 10.1101/2024.09.04.611236

**Authors:** Biana Bernshtein, Julia A. Zhiteneva, Jeshina Janardhanan, Chanchal Wagh, Meagan Kelly, Smriti Verma, Wonyeong Jung, Salima Raiyan Basher, Mohammad Ashraful Amin, Shakil Mahamud, Nazmul Hasan Rajib, Fahima Chowdhury, Ashraful Islam Khan, Richelle C. Charles, Peng Xu, Pavol Kováč, Subhra Chakraborty, Robert W. Kaminski, Galit Alter, Taufiqur R. Bhuiyan, Firdausi Qadri, Edward T. Ryan

## Abstract

Shigellosis is the second leading cause of diarrheal death in children younger than five years of age globally. At present, there is no broadly licensed vaccine against shigella infection. Previous vaccine candidates have failed at providing protection for young children in endemic settings. Improved understanding of correlates of protection against Shigella infection and severe shigellosis in young children living in endemic settings is needed. Here, we applied a functional antibody profiling approach to define Shigella-specific antibody responses in young children versus older individuals with culture-confirmed shigellosis in Bangladesh, a Shigella endemic area. We analyzed Shigella-specific antibody isotypes, FcR binding and antibody-mediated innate immune cell activation in longitudinal serum samples collected at clinical presentation and up to 1 year later. We found that higher initial Shigella O-specific polysaccharide (OSP)-specific and protein-specific IgG and FcγR binding levels correlated with less severe disease regardless of patient age, but that individuals under 5 years of age developed a less prominent class switched, FcR-binding, functional and durable antibody response against both OSP and protein Shigella antigens than older individuals. Focusing on the largest cohort, we found that functional *S. flexneri* 2a OSP-specific responses were significantly induced only in individuals over age 5 years, and that these responses promoted monocyte phagocytosis and activation. Our findings suggest that in a Shigella endemic region, young children with shigellosis harbor a functional antibody response that fails to maximally activate monocytes; such a response may be important in facilitating subsequent innate cell clearance of Shigella, especially via recruitment and activation of polymorphonuclear cells capable of directly killing Shigella.

## Introduction

Diarrheal diseases are a major cause of morbidity and mortality in young children living in low-resource settings^1^. Among bacterial and viral pathogens causing moderate to severe diarrhea, Shigella is the second leading cause in children ages 1-2 years of age and the leading cause in children 3-5 years of age^2^. Yet, our understanding of the correlates of protection against severe diarrheal disease caused by Shigella are cursory^3^, especially in young children living in endemic settings. Young children harbor an immune system distinct from adults, partly due to accumulated exposure to pathogens that shapes immune responses and immunological memory over time in adults^4^. As young children under 5 years of age are the main target population for a vaccine against Shigella, understanding the unique protective immune responses to shigellosis in young children living in Shigella-endemic settings is critical. *Shigella* species and serotypes share several antigens (such as invasion protein antigens: Ipa), but also have distinct O-specific polysaccharide (OSP) components of lipopolysaccharide (LPS). Protection against shigellosis is serotype-specific, although immune responses to shared antigens may also contribute to protection^5,6^. Previous studies have found a protective role for LPS-specific IgG against Shigella infection in adults living in endemic settings^7,8^, and we have recently shown that OSP-specific functional IgA protects adults against incident Shigella infection in a high Shigella burden setting in Peru^9^ . A number of shigella vaccines are in development, most targeting at least in part the most common Shigella OSPs^10–12^. Previously an OSP-conjugate vaccine showed efficacy that decreased with age, with very low efficacy in children ages 1-3 years^13^. As vaccines against Shigella move forward in development and evaluation, it will thus be important to better understand age-dependent differences in immune responses and protection against disease severity, especially in young children living in Shigella-endemic areas.

Accordingly, we analyzed longitudinal functional antibody profiles of children and adults living in Bangladesh who presented for clinical care with stool culture-confirmed shigellosis caused by one of 4 prevalent species/strains of Shigella: *S. flexneri* 2a, 3a, 6 and *S. sonnei*. We characterized serum antibody antigen-specific, FcR-binding, and functional antibody responses including innate cell activation from time of presentation for clinical care until 1-year post infection, comparing responses in young children versus older individuals in this highly-endemic location and in which shigella vaccines may be tested in phase III studies.

## Results

### Dynamics of immune responses to shigellosis

We analyzed antibody profiles in patient serum presenting with stool culture-confirmed Shigella infection with one of 4 different species or strains of Shigella: *Shigella flexneri* 2a (n=20), 3a (n=11), 6 (n=5) or *Shigella sonnei* (n=19) on days 2, 7, 30, 90, 180 and 360 after presenting for clinical care (**SuppTable 1**). With regard to pan-anti-Shigella Ipa protein responses, we found significant rises in Shigella protein-specific IgG responses against IpaB, IpaC and IpaD (**Supp Fig 1**) that persisted in circulation for up to 90 days (for IpaB) or 30 days (for IpaC and IpaD) after clinical presentation. Shigella protein-specific IgG induced by infection binds to FcγRs; however, only IpaB-specific antibodies binding to FcγR3b on day 7 and day 30 reached statistically significant induction levels (**Supp Fig 1**). Shigella protein-specific IgA and FcαR binding were, in general, not significantly induced, although there was a significant decrease in IpaB-specific FcαR binding levels at day 30 compared to day 7 (**Supp Fig 1**). We also analyzed OSP-specific responses in patients infected with either *S. flexneri* 2a or *S. sonnei*, the two most prominent strains of Shigella world-wide and the largest groups in the cohort, to better focus analysis of OSP-specific responses since they vary by species and serotype (**Fig 1**). While IgG responses against *S. sonnei* OSP were significantly induced by day 7 after presentation and remained significantly elevated on day 30, *S. flexneri* 2a OSP-specific IgG responses also increased after initial sampling but did not reach statistical significance (**Fig 1A**). By day 90 after clinical presentation, *S. sonnei* OSP-specific IgG levels were comparable to levels at the time of initial clinical presentation (**Fig 1A**). *S. sonnei* OSP-specific IgG responses induced by infection were dominated by IgG1 and IgG2, and while FcγR binding levels were elevated on day 7 and day 30 compared to day 2, they did not reach statistical significance, possibly due to small sample number (**Fig 1A**). Additionally, while IgA1 responses against *S. sonnei* OSP did not change over time, binding to FcαR was significantly induced on day 7 following clinical presentation, although it returned to the baseline level on day 30, suggesting a change in functionality of OSP-specific IgA (**Fig 1A**). Although there was no significant change in *S. flexneri* 2a OSP-specific responses when considering the entire cohort of *S. flexneri* 2a infected individuals, there was a significant increase in OSP-specific IgG and FcγR2a binding responses on days 7 and 30 in individuals over 5 years of age, and not in younger children (**Fig 1B**).

**Fig 1.**
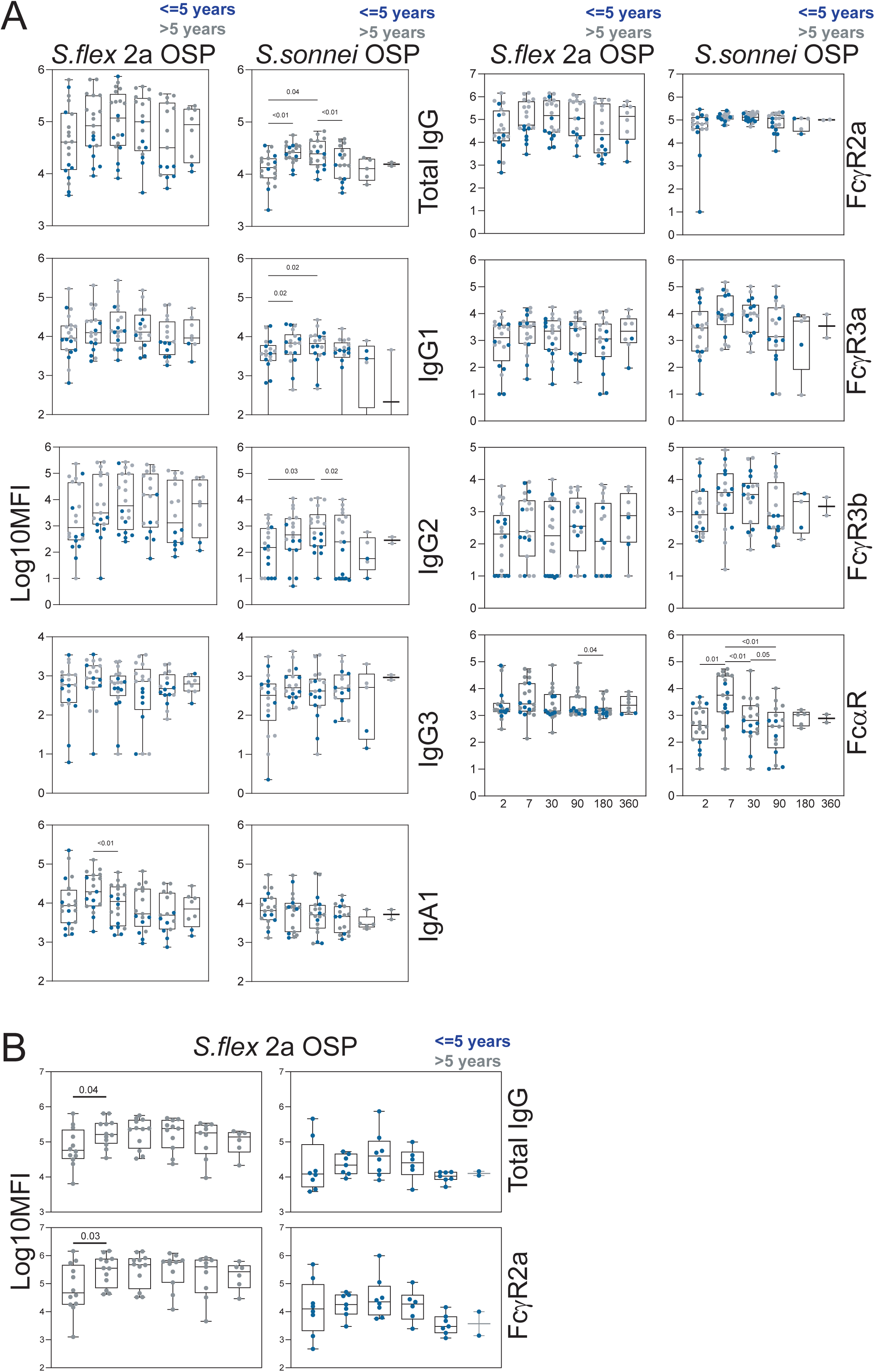
Shigella O-specific polysaccharide (OSP) antibody responses to shigellosis. **(A)** *S. flexneri* 2a and *S. sonnei* OSP-specific total IgG, IgG1, IgG2, IgG3, IgA1 and binding to FcγR2a, FcγR3a, FcγR3b and FcαR in serum of individuals presenting with corresponding strain stool culture-confirmed shigellosis in Bangladesh. **(B)** *S. flexneri* 2a OSP-specific total IgG and FcγR2a binding in individuals younger or older than 5 years of age. (p-value calculated by Mixed effects analysis followed by Holm-Sidak’s multiple comparisons test)

### Shigella-specific antibody features associated with disease severity

Our understanding of correlates of protection against severe shigellosis in endemic settings is incomplete. We sought to find antibody features associated with less severe disease in this cohort of individuals who presented for clinical care with stool culture-confirmed Shigella infection in Bangladesh. We calculated a disease severity index (DSI) for each patient encompassing information collected upon presentation, considering the presence of blood in stool, fever, vomiting and number of loose stools per day^14^. Next, we correlated the calculated DSI of each patient with Shigella protein- and OSP-specific antibody features on day 2 after clinical presentation (**Fig 2**). To compile all confirmed Shigella cases across 4 different species and strains, we used OSP responses corresponding to the infection strain for each subject (infection strain OSP). Overall, antibody levels on day 2 were inversely correlated with disease severity, suggesting that Shigella-specific antibodies may be protective against severe shigellosis in endemic settings (**Fig 2**). Specifically, IpaB- and IpaC-specific IgG and FcγR binding levels significantly inversely correlated with shigellosis severity (**Fig 2**). However, IpaD-specific antibodies were not significantly correlated with risk or protection against shigellosis (**Fig 2**). OSP-specific IgG and binding to FcγR2a and FcγR3a were also significantly correlated with protection (**Fig 2**). Of note, both protein- and OSP-specific IgA, IgM and FcαR binding were not significantly correlated with disease severity, though for FcαR binding there was a trend of inverse correlation, suggesting some level of protection (**Fig 2**). OSP-specific antibody mediated protection against shigellosis is largely strain specific^7,15,16^; however, there is evidence for potential cross-protection between different species and strains, especially relating to shared antigens such as Ipa proteins ^5,6,17^. To test for possible OSP-specific antibody associated cross-protection in the cohort, we also correlated antibody responses against *S. flexneri* 2a, 3a and *S. sonnei* OSP on day 2 with DSI of patients infected with heterologous strains (**Supp Fig 2**). We found significant inverse correlation between *S. sonnei* OSP-specific total IgG, IgG2 and FcγR2a levels and DSI in patients infected with *S. flexneri* 2a on day 2, suggesting that *S. sonnei* OSP-specific IgG may correlate with protection against severe shigellosis caused by *S. flexneri* 2a. We did not find significant correlation for the reverse; namely *S. flexneri* 2a OSP-specific IgG levels did not correlate with protection against severe shigellosis caused by *S. sonnei,* nor did we find any correlation between DSI and OSP-specific antibody levels for other strains (**Supp Fig 2**). To assess whether the inverse correlation between antibody features and DSI was skewed by time from infection, we analyzed a subset of samples collected from patients enrolled in the study no later than 48hrs after symptom onset (n=23) (**Supp Fig 3**). Overall, we found similar inverse correlation in this subset of patients as in the full cohort, although some correlations did not reach statistical significance, possibly due to the smaller number of samples (**Supp Fig 3**).

**Fig 2.**
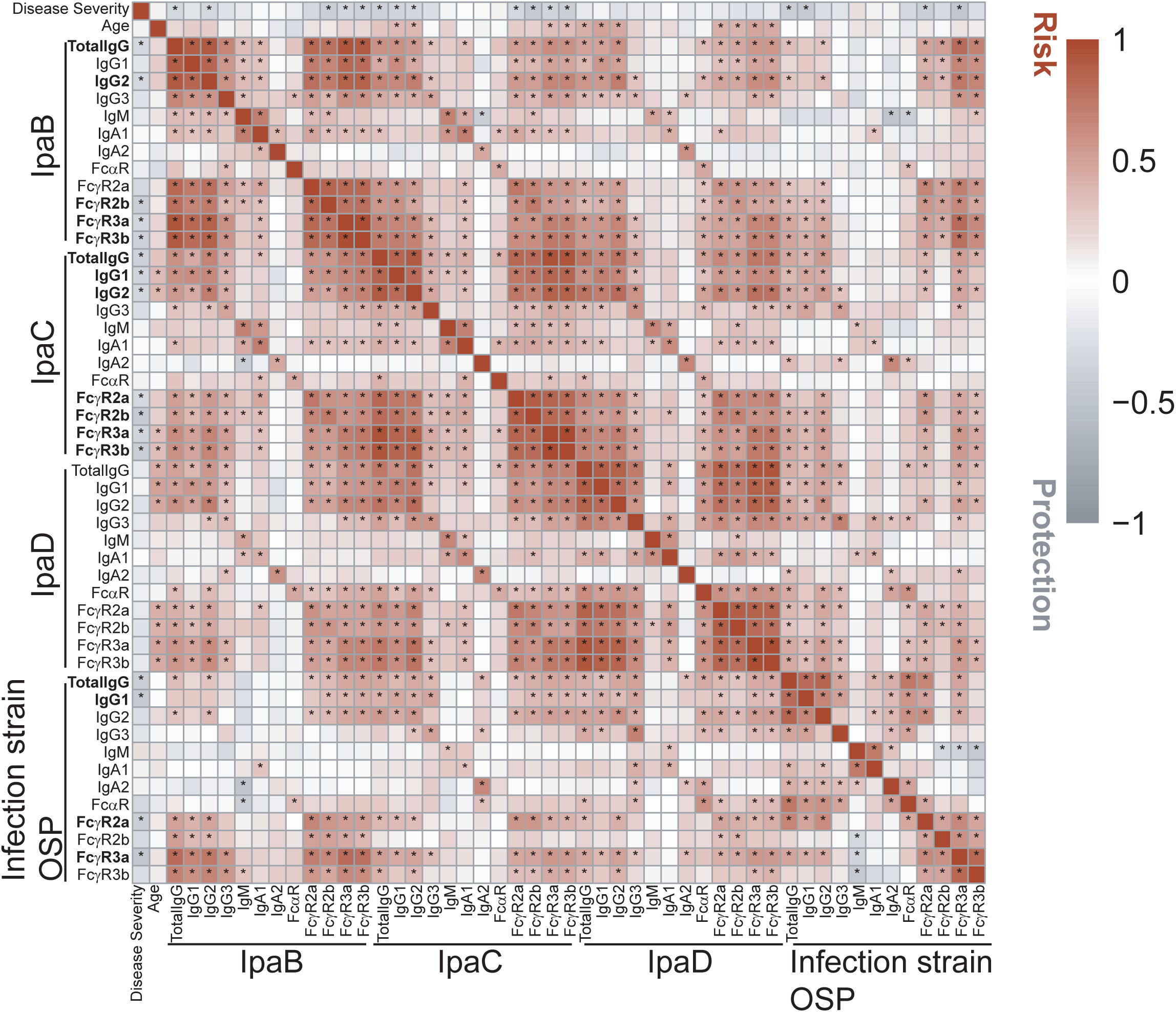
Impact of antibody profiles on disease severity. Heat map of Spearman correlation of disease severity and age with Shigella-specific antibody features in serum of individuals presenting with stool culture-confirmed shigellosis on day 2 of clinical care. (*Benjamini-Hochberg corrected p-value<0.05)

As FcγR binding levels were significantly associated with protection, we sought to determine correlations between antibody-mediated innate immune cell effector functions and DSI (**Fig 3**). In this analysis we focused on serum samples collected on day 2 from individuals infected with *S. flexneri* 2a or *S. sonnei*, the most prevalent strains causing shigellosis in the cohort. Antibody dependent neutrophil phagocytosis (ADNP) mediated by IpaB-specific antibodies significantly inversely correlated with DSI, suggesting that IpaB-specific functional antibodies engage and activate neutrophils that protect against severe shigellosis (**Fig 3A**). Importantly, IpaB-specific ADNP on day 2 did not significantly correlate with patients’ age (**Fig 3A**). IpaC- and infection strain OSP-specific ADNP did not significantly correlate with disease severity (**Fig 3A**). Antibody dependent cellular phagocytosis (ADCP) mediated by IpaB, IpaC or infection strain OSP-specific antibodies did not significantly correlate with disease severity; however, IpaB ADCP showed a trend of inverse correlation with disease severity (**Fig 3B**). IpaB and infection strain OSP-specific antibody dependent monocyte phagocytosis (ADMP) did not correlate with disease severity (**Fig 3C**). Antibody dependent complement deposition (ADCD) mediated by IpaB, IpaC, IpaD and infection strain OSP-specific antibodies did not significantly correlate with protection (**Fig 3D**). IpaB and infection strain OSP-specific ADCD showed a trend of positive correlation with DSI, suggesting that complement deposition mediated by IpaB and OSP-specific antibodies on day 2 associate with severe disease (**Fig 3D**). In general, DSI did not significantly correlate with age of patients (**Fig 2, 3A-D**), and comparing DSI of individuals older and younger than 5 years of age showed no significant difference (**Fig 3E**). Overall, inverse correlations of antibody features on day 2 after clinical presentation with DSI suggest that Shigella-specific antibodies are protective regardless of age.

**Fig 3.**
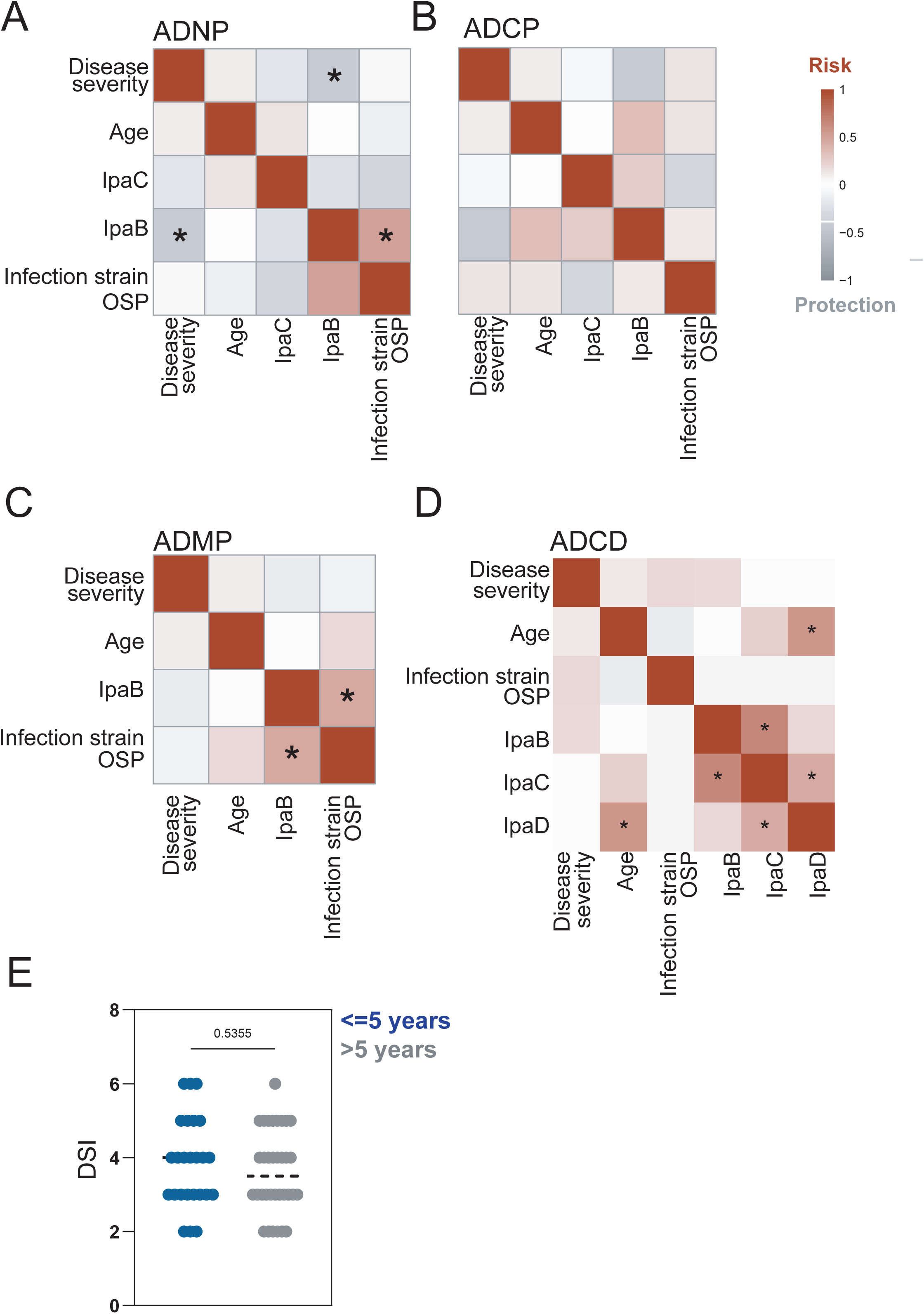
Impact of functional antibodies on disease severity. Heat map of Spearman correlation of disease severity and age with Shigella-specific ADNP **(A)** ADCP **(B)** ADMP **(C)** and ADCD **(D)** in serum of individuals presenting with stool culture-confirmed shigellosis on day 2 after clinical presentation. (*Benjamini-Hochberg corrected p-value<0.05). **(E)** Disease severity index (DSI) of individuals younger or older than 5 years of age.

### Age-dependent response to Shigella infection

Children under 5 years of age bear a very large global burden of shigellosis^18–20^. Thus, we sought to interrogate Shigella-specific immune responses in individuals over 5 years of age versus younger children (**Fig 4**). We applied a non-biased machine learning approach to define antibody features enriched in individuals older or younger than 5 years of age in *S. flexneri* 2a (**Fig 4A-B**) and *S. sonnei* (**Supp Fig 4A-B**) infected patients on day 7 after clinical presentation. Following a Least Absolute Shrinkage and Selection Operator (LASSO) down selection of features, a Partial Least Squares-Discriminant Analysis (PLS-DA) was used to visualize the data (**Fig 4, SuppFig4**). Out of total 142 antibody features measured, 7 were down selected by LASSO as sufficient to separate the groups in *S. flexneri* 2a infected individuals (**Fig 4A**). OSP-specific total IgG and IpaD-specific IgG1 and IgG2 were enriched in older individuals compared to children younger than 5 years of age (**Fig 4A**). Importantly, as LASSO eliminates co-correlated features, we built a correlation network of selected features with other antibody features (Pearson co-efficient>0.07, adjusted p value<0.01) (**Fig 4B**). Co-correlates network of selected features of *S. flexneri* 2a infected individuals on day 7 revealed correlation of OSP-specific total IgG with OSP-specific IgG2 and binding of OSP-specific antibodies to FcγR2a (**Fig 4B**). IpaD-specific IgG2 represents a larger network of co-correlated IpaD-specific IgG that bind to FcγRs and IpaD-specific IgA enriched in older individuals (**Fig 4B**). IpaB ADCP was enriched in older individuals while IpaB-specific IgA1 and IgA2 were enriched in individuals under age 5. Additionally, IpaB-specific IgA1 correlated with IpaB-specific total IgG and IgG1 and binding of IpaB-specific antibodies to FcγR2a (**Fig 4B**). IpaB-specific IgA1 correlated with *S. sonnei* OSP-specific total IgG while IpaB-specific IgA2 correlated with *S. sonnei* OSP-specific IgA2 (**Fig 4B**). Finally, *S. flexneri* 2a OSP ADMP was enriched in older individuals (**Fig 4A**).

**Fig 4.**
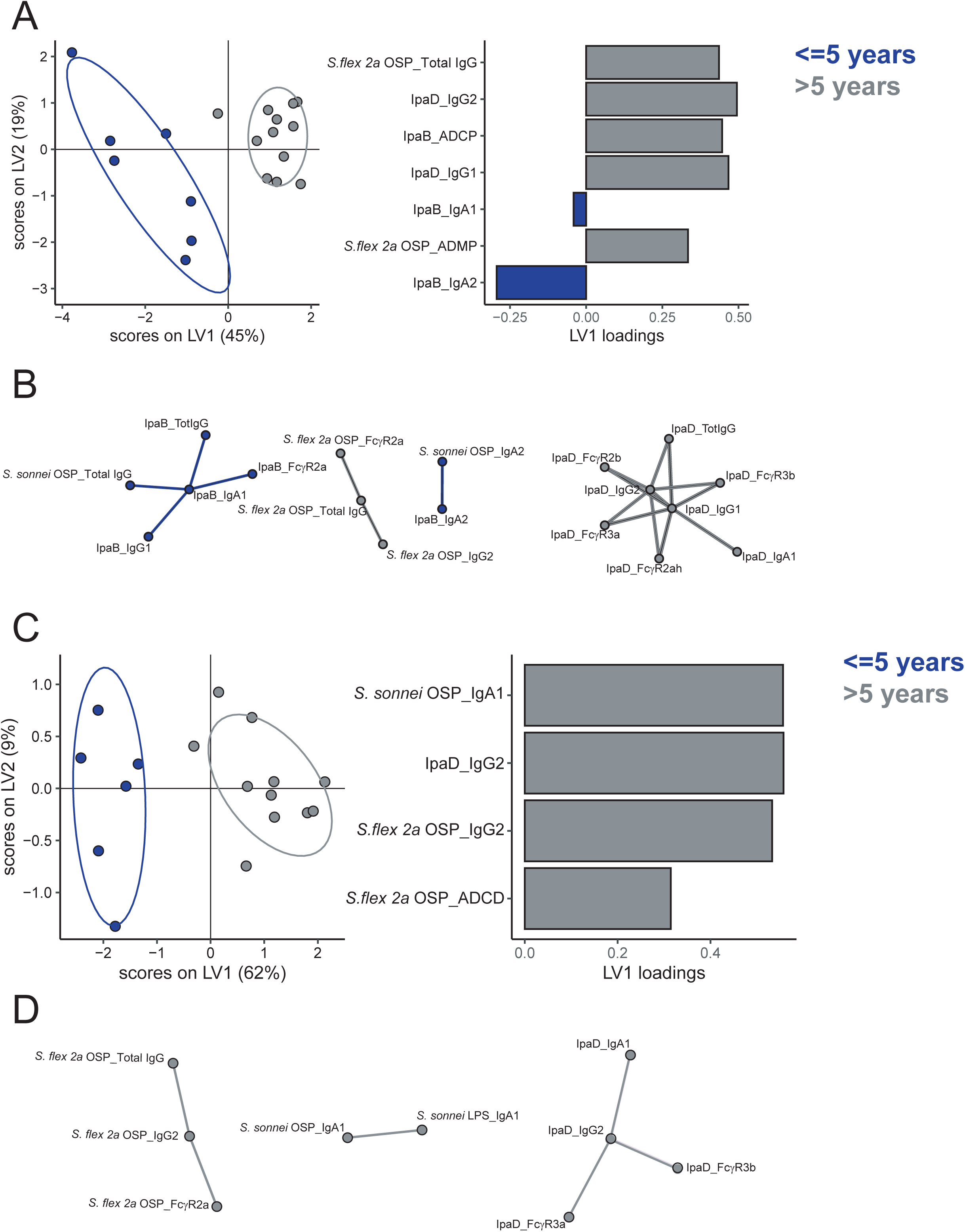
Age-dependent differences in immune response to shigellosis. Score plots and LV1 loadings of PLS-DA model using LASSO selected features of Shigella-specific antibody isotypes and FcR binding in serum of individuals on day 7 **(A-B)** or day 90 (**C-D**) after clinical presentation with stool culture-confirmed shigellosis caused by *S. flexneri* 2a and corresponding correlation networks of selected antibody features (**B,D**) (Spearman rank coefficient R>0.7, Benjamini-Hochberg corrected p<0.01)

We applied the same analysis pipeline to separate individuals younger and older than 5 years of age infected with *S. sonnei* on day 7 after clinical presentation (**Supp Fig 4A-B**). LASSO selected 2 out of 142 antibody features analyzed to separate the two groups, both were enriched in older individuals (**Supp Fig 4A**). IpaD-specific FcγR binding antibodies were enriched in older *S. sonnei* infected individuals as well and were part of a broader network of IpaD-specific and *S. flexneri* 2a OSP-specific antibodies enriched in older individuals (**Supp Fig 4A-B**). Additionally, *S. sonnei* OSP-specific IgG2 were enriched in older individuals infected with *S. sonnei (***Supp Fig 4A**). Overall, infection strain OSP-specific IgG responses were enriched in older individuals in both *S. flexneri* 2a and *S. sonnei* infected individuals on day 7 (**Fig 4A-B**, SuppFig4A-B).

To determine long-term differences between antibody responses in individuals over and under 5 years of age, we applied the same analysis pipeline to samples collected 90 days after clinical presentation (**Fig 4C-D, Supp Fig 4C-D**). For *S. flexneri* 2a infected individuals on day 90, LASSO selected 4 out of 142 measured antibody features that were enriched in individuals over age 5 (**Fig 4C**). *S. sonnei* OSP-specific IgA1, IpaD-specific IgG2 and *S. flexneri* 2a OSP-specific IgG2 and ADCD were enriched in individuals over 5 years of age on day 90 after presentation (**Fig 4C**). *S. flexneri* 2a OSP-specific IgG2 correlated with total IgG and binding to FcγR2a, while IpaD-specific IgG2 correlated with IgA1 and binding to FcγR3a and FcγR3b (Pearson co-efficient>0.07, adjusted p value<0.01) (**Fig 4D**). For *S. sonnei* infected individuals on day 90, LASSO selected 3 features to separate individuals 5 years of age versus younger – *S. sonnei* LPS-specific total IgG, OSP-specific IgG2 and IpaC-specific antibody binding to FcγR2a (**Supp Fig 4C**). Co-correlates network of these selected features revealed a broad and functional antibody network present in individuals over age 5 years on day 90 after presentation (**Supp Fig 4D**).

Given these robust age-dependent differences in antibody responses and enrichment of OSP-specific antibodies in individuals over age 5 years, we next focused on individuals infected with *S. flexneri* 2a to determine temporal dynamics of *S. flexneri* 2a OSP-specific antibodies in older and younger individuals (**Fig 5A**). *S. flexneri 2a* OSP-specific total IgG and FcγR2a-binding were elevated in older individuals at day 2, and remained robustly elevated over time, as long as 1 year after clinical presentation (**Fig 5A**). Binding to FcγR3a and FcγR3b of OSP-specific antibodies was also elevated in older individuals over time; however, that difference did not reach statistical significance (**Fig 5A**). OSP-specific IgA1 levels were significantly elevated in older individuals on day 30 after clinical presentation (**Fig 5B**). Interestingly, OSP-specific IgA2 showed the opposite trend, as it was elevated on day 30 in individuals under age 5 years (**Fig 5B**). FcαR binding of OSP-specific antibodies was slightly elevated in older individuals on day 7 (**Fig 5B**). Finally, OSP-specific ADMP was enriched in older individuals over time as well, while ADNP and ADCD were not different between younger and older individuals (**Fig 5B**).

**Fig 5.**
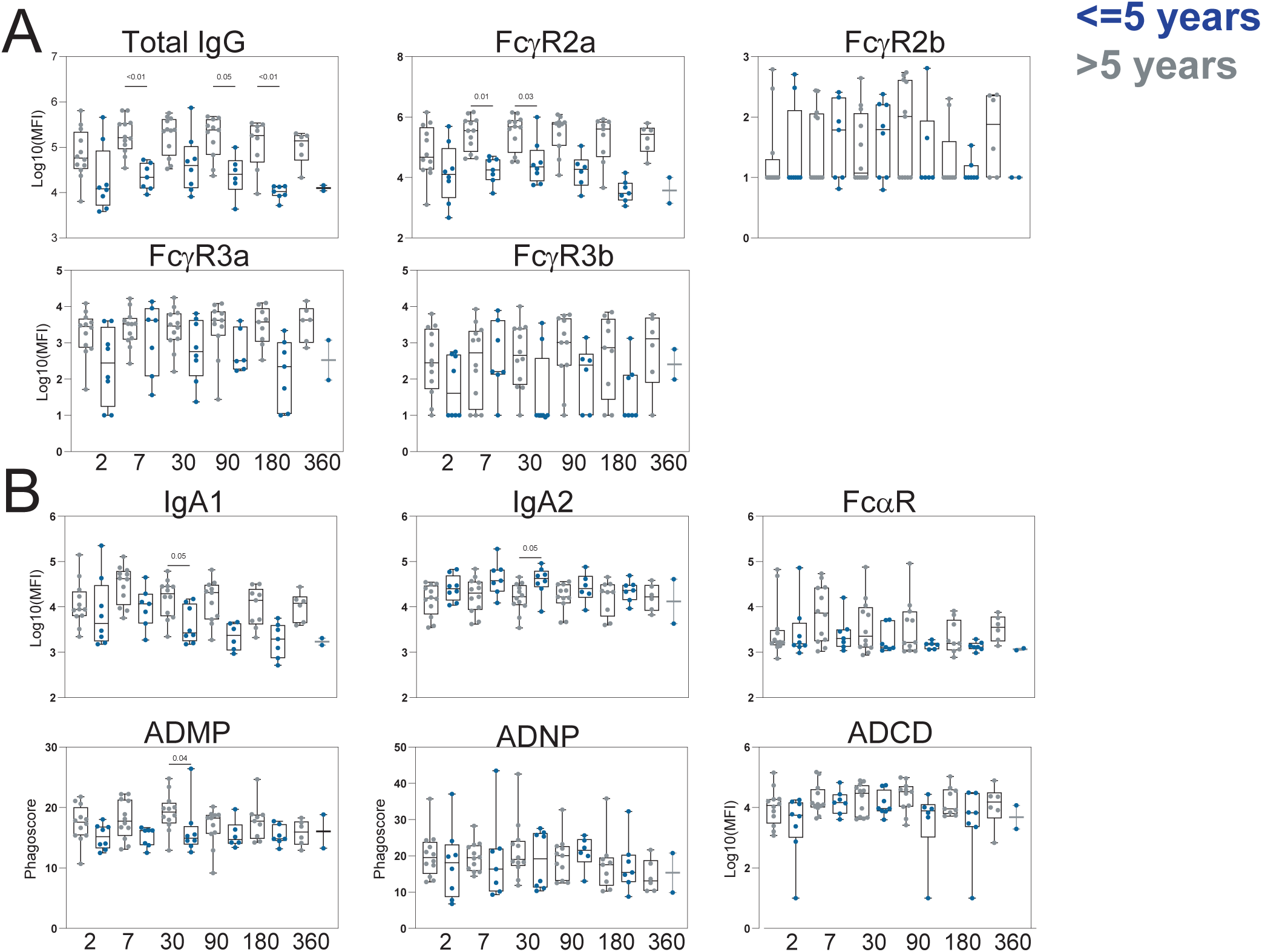
Dynamics of age-dependent longitudinal responses to S. flexneri 2a OSP. *S. flexneri* 2a OSP-specific **(A)** total IgG and binding to FcγR2a, FcγR2a, FcγR3a and FcγR3b, IgA1, IgA2, FcαR, ADMP, ADNP and ADCD in serum of individuals over and under age 5 presenting with stool culture-confirmed *S. flexneri* 2a shigellosis in Bangladesh. (p-value calculated by Kruskal-Wallis test)

Given elevated levels of both OSP-specific IgG and IgA1 with increased FcγR and FcαR binding in older individuals at early time points after clinical presentation, together with higher OSP ADMP in these individuals, we sought to determine the contribution of IgA and FcαR binding versus IgG and FcγR binding to antibody mediated monocyte phagocytosis and activation. To address this question, we specifically and efficiently depleted IgG or IgA from serum of individuals over age 5 collected at days 2, 7 and 30 after presentation (**Fig 6A**). OSP ADMP was significantly reduced by IgG depletion, but not by IgA depletion, suggesting that OSP-specific IgG and FcγR binding facilitate monocyte phagocytosis (**Fig 6B**). As monocytes play a critical role in Shigella pathogenesis^21–23^, we sought to deeply investigate OSP-specific IgG mediated activation of monocytes^24^. We depleted IgG from serum samples of individuals > 5 years of age collected on days 2, 7, and 30 after presentation and measured antibody-mediated monocyte phagocytosis and induction of monocyte activation markers including CD16, CD86, CCR2, MHCII and CX3CR1. OSP ADMP was significantly reduced following IgG depletion for serum collected on days 2, 7 and 30 after clinical presentation (**Fig 6C**), as well as surface expression levels of the co-stimulatory molecule CD86 and activation markers CD16 and CX3CR1 (**Fig 6C**). Levels of the chemokine receptor CCR2 and antigen presentation complex MHCII expression on the surface of monocytes were downregulated by OSP-specific IgG, significantly in serum collected on days 2 and 7 after clinical presentation (**Fig 6C**). These data suggest that OSP-specific IgG, elevated predominantly in *S. flexneri* 2a infected adults but not in young children, binds to Fcγ receptors expressed on the surface of monocytes to facilitate phagocytosis, expression of costimulatory molecules and activation markers, and reduce expression levels of CCR2 and MHCII.

**Fig 6.**
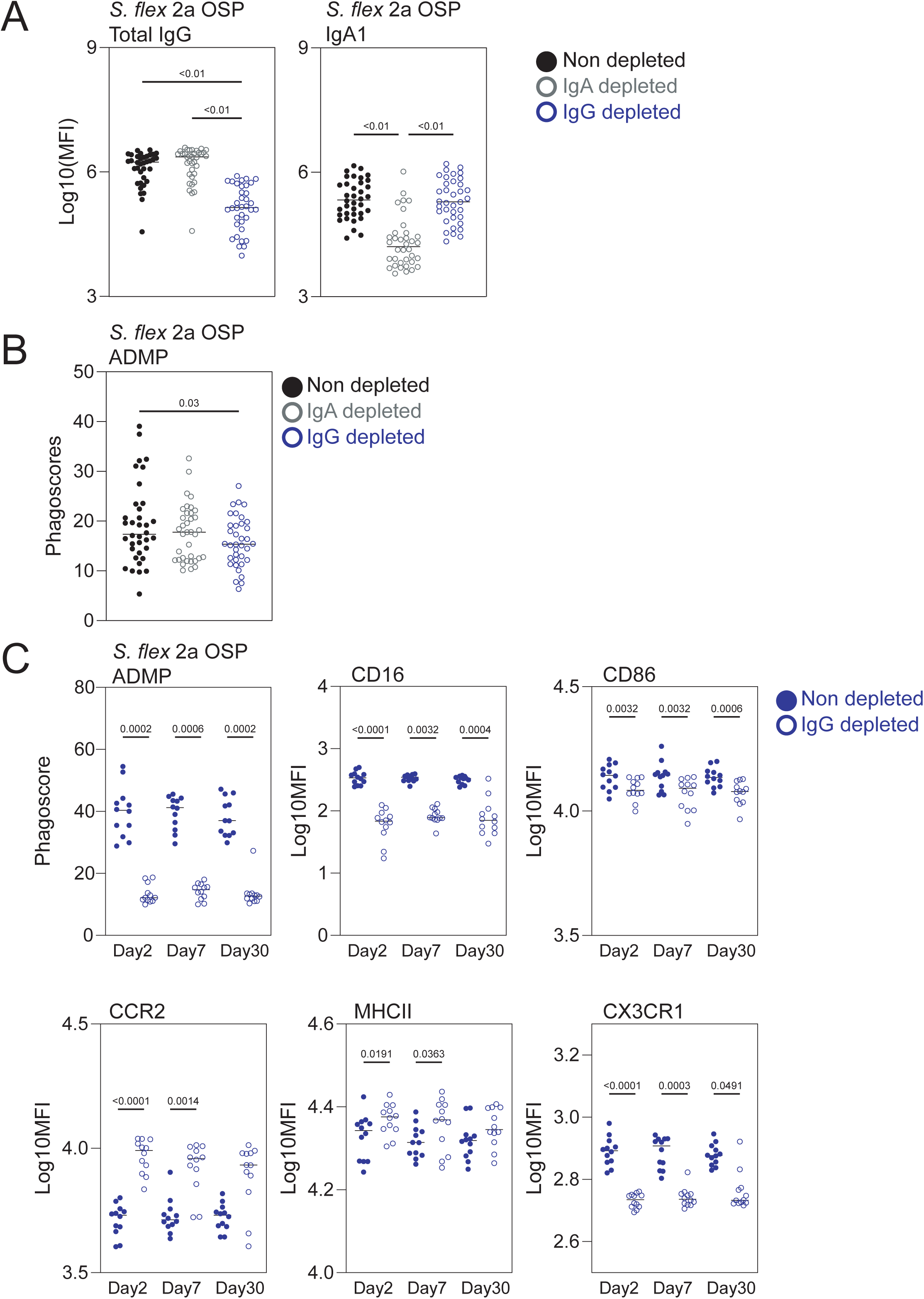
S. flexneri 2a OSP-specific IgG activates monocytes. **(A)** *S. flexneri* 2a OSP-specific IgG and IgA1 levels in non-depleted, IgG-depleted and IgA-depleted serum from adults presenting with stool culture-confirmed infection with *S. flexneri* 2a on days 2, 7 and 30 after clinical presentation **(B)** *S. flexneri* 2a OSP-specific ADMP in non-depleted, IgA-depleted or IgG-depleted serum of adults infected with *S. flexneri* 2a on days 2, 7 and 30 after clinical presentation **(C)** *S. flexneri* 2a OSP-specific ADMP, CD16, CD86, CCR2, MHCII and CX3CR1 surface expression levels of monocytes stimulated with *S. flexneri* 2a OSP-specific immune complexes of non-depleted or IgG-depleted serum of adults infected with *S. flexneri* 2a on days 2, 7 and 30 after clinical presentation. (p-value calculated with Friedman test)

## Discussion

Individuals living in Shigella-endemic settings are exposed throughout their lives to repetitive infections, stimulating immune responses and memory that provide protection against disease. In early life, following waning of maternally transferred immunity and prior to sufficient exposure, Shigella infection is highly prevalent and results in severe outcomes^1^. An efficient Shigella vaccine capable of eliciting protective immune response in young children living in endemic settings is critically needed. Here, we found significant differences in immune responses to Shigella infection between individuals over and under the age of 5 years living in a Shigella-endemic setting. On day 7 after clinical presentation, individuals over 5 years of age harbored a class switched, FcR-binding, functional and durable antibody response against both OSP and protein Shigella antigens. As OSP-specific antibodies are an established correlate of protection against shigellosis^3,7,8,16^ and in light of the fact that ongoing attempts to develop Shigella vaccines are often based on OSP antigens^25–27^, we focused on age-dependent dynamics and functionality of OSP-specific antibodies in patients infected with the most common cause of shigella infection globally and in Bangladesh, *S. flexneri* 2a. We found that individuals over 5 years of age infected with *S. flexneri* 2a responded to infection by inducing functional OSP-specific IgG that binds to FcγRs and promotes monocyte phagocytosis and activation. Shigella pathogenesis is intimately involved in host cell interactions. Nonactivated macrophages are unable to kill Shigella directly^23,28,29^. Once ingested by nonactivated macrophages, Shigella escape from the phagosome and initiate an inflammatory cascade resulting in a pyroptotic death of the macrophage^23^. Cell death results in release of the bacteria, facilitating invasion of the target colonic host cell; however, it also results in recruitment of polymorphonuclear cells that are more resistant to Shigella-induced killing and that are eventually capable of directly killing Shigella^23,30^. This killing by polymorphonuclear cells is also facilitated in the presence of anti-Shigella antibodies that enhance bacterial opsonization^30–33^. Our data thus suggest that OSP-specific IgG responses facilitate monocyte phagocytosis and activation that may lead to improved polymorphonuclear cell recruitment and Shigella killing. Our findings that individuals under age 5 years of age harbor lower and less durable levels of OSP-specific IgG, as well as reduced FcγR binding of OSP-specific antibodies and OSP ADMP, may thus suggest a possible contributor to susceptibility to severe shigellosis with its associated morbidity and mortality in young children in endemic areas.

We previously found that OSP-specific FcαR binding antibodies capable of activating polymorphonuclear cells are protective against incident Shigella infection in a high burden setting in an endemic adult population in Peru^9^. Since our study in Bangladesh was designed to evaluate individuals already infected with Shigella, we were unable to ascertain protection from infection in this current study. However, our design did allow us to assess correlation of immune responses with disease severity once infected. Our two analyses suggest two layers of OSP-specific antibody mediated protection against shigellosis: one involved in preventing infection, mediated by FcαR binding IgA that activates bactericidal activities of neutrophils, potentially in the intestinal lumen; and the other mediated by IgG that maximally activates monocytes to facilitate innate cell handling of Shigella, possibly through facilitating increased recruitment of polymorphonuclear cells within the intestinal lamina propria following bacterial invasion. In our current study, we also found that FcγR binding IpaB-specific responses that activate ADNP were associated with protection from severe disease, further underscoring the role of polymorphonuclear cells in mediating protection.

Importantly, disease severity in our cohort was not significantly different between infected individuals over and under 5 years of age. We found that Shigella-specific antibody levels and FcR-binding on day 2 after clinical presentation inversely correlated with disease severity, independent of age, suggesting that functional antibodies are overall protective against severe disease. However, as OSP-specific IgG and FcγR binding were correlated with protection, and more durable in individuals over 5 years of age, this suggests that these individuals may be better and longer protected against future infections, perhaps by consolidating immune responses primed by previous exposures. Further, our data suggest IgG-mediated activation of monocytes as a mechanism of action for OSP-specific protective antibodies.

Interestingly, in a sub-analysis we also found that *S. sonnei* OSP-specific antibodies correlated with protection against severe disease in *S. flexneri 2a* infected patients. Since the OSPs of these two species are distinct, this possible cross-species correlation could be mediated by antibodies binding to the shared core oligosaccharide connecting OSP to lipid A or could represent a marker of previous infection inducing protection mediated by other arms of the immune system, for example T cells. This potential cross-protection could facilitate Shigella vaccine development and should be further tested in controlled infection models. In addition to OSP-specific antibodies, IpaD-specific antibody levels were positively correlated with age in our cohort. This could point to IpaD-specific responses as markers of previous exposures and should be tested further in sero-surveillance studies.

Our study has several limitations. First, since we are collecting samples in endemic settings from individuals naturally infected with Shigella, we were limited by the number of samples available and the distribution of Shigella strains and volunteer ages. This limited analysis of age-dependent antibody responses to *S. flexneri* 2a and *S. sonnei*. Future analysis will focus on other Shigella serotypes. Our analysis was also limited to individuals seeking medical attention, hence we could not study protection against infection itself as all of the individuals in our cohort were infected, nor could we control for milder disease not prompting seeking of facility-based medical care. Our study was also unable to distinguish whether the observed differences in immune responses reflected differences in age-dependent immune handling or exposure history in this highly endemic area. We also used a disease severity score (DSI) based on a scoring method used in human challenge studies^14^, one specific for young children with shigella infection in an endemic area has yet to be developed. Lastly, our functional cellular-based assays are bead-based, hence we were not able to differentiate effects of OSP-specific antibodies on bacterial infection of monocytes, including activation of pyroptotic pathways by living Shigella in this study, but we plan to evaluate this aspect separately.

In summary, we found that higher initial OSP-specific and protein-specific IgG and FcγR binding levels correlated with less severe disease regardless of patient age, but that individuals under age 5 years developed a less pronounced class switched, FcR-binding, functional and durable antibody response against both OSP and protein Shigella antigens than older individuals. Focusing on the largest cohort, we found that functional *S. flexneri* 2a OSP-specific responses were only significantly induced in individuals over age 5 years in this highly endemic area, and that these responses promoted monocyte phagocytosis and activation. Our findings suggest that young children with shigellosis are less able to mount a functional antibody response able to activate monocytes as compared to older individuals in a Shigella endemic region; such a response may be important in facilitating subsequent innate cell clearance of Shigella, especially via recruitment and activation of polymorphonuclear cells capable of directly killing Shigella. Vaccines capable of inducing potent immune responses with a functional component in children younger than 5 years of age, such as the ability to activate monocytes and polymorphonuclear cells, may be required to protect this most vulnerable population from shigellosis.

## Methods

### Study design and participants

Following an informed consent process and enrollment of eligible patients, blood was collected via venipuncture from study participants on the second day following clinical presentation to the International Centre for Diarrhoeal Disease Research in Dhaka, Bangladesh (icddr.b) (**SuppTable 1**). Patients were screened by a rapid Shigella diagnostic assay RLDT^34,35^, and study enrollees had microbiologic isolation of *S. flexneri* 2a, 3a, 6 or *S. sonnei* from their stool. Stool was plated on MacConkey agar and Salmonella Shigella agar (SS) for isolating *Shigella* bacteria. Serotype was confirmed by anti-sera agglutination test (MAST Assure, UK). Blood was collected at time of study enrollment (day 2; the day after clinical presentation and study enrollment), then on day 7, 30, 90, 180 and 360. Sera was isolated from blood and frozen for subsequent analysis. Patients greater than 60 months of age were classified as older than 5 years of age. Disease severity index (DSI) was calculated as previously described^6,14^, namely individuals received 1 point for presence of either of the following at presentation: blood in diarrhea, fever or vomiting; number of stools per 24 hours was categorized into groups: 1 or more as 1 point, 3 or more as 2 points, 8 or more is 3 points. Points were then added to calculate a DSI for every patient (**SuppTable 1**).

### Procedures

#### Luminex

*Shigella*-specific antibody subclass/isotype and Fcγ-receptor (FcγR) binding levels were assessed using a 384-well based customized multiplexed Luminex assay, as previously described^36^. IpaB (WRAIR), IpaC (WRAIR), IpaD (WRAIR), *S. flexneri* 2a 2457T LPS (WRAIR), *S. flexneri* 3a strain J17B LPS (WRAIR), *S. flexneri* 6 Sf6_1114_19 LPS (icddrb), *S. sonnei* Moseley LPS (WRAIR), *S. flexneri* 2a OSP, *S. flexneri* 3a OSP, *S. flexneri* 6 OSP and *S. sonnei* OSP were used to profile -specific humoral immune response. OSP was purified from *S. flexneri* 2a Sf2a260214_1 LPS (icddrb), *S. flexneri* 3a Sf3a050214_5 LPS (icddrb), *S. flexneri* 6 Sf6_1114_19 LPS and *S. sonnei* Moseley LPS by acid hydrolysis and size exclusion chromatography and conjugated to BSA^37–42^ . Tetanus toxin and Ebola (CEFTA, Mabtech Inc) were used as control antigens. Protein antigens were coupled to magnetic Luminex beads (Luminex Corp) by carbodiimide-NHS ester-coupling (Thermo Fisher). OSP and LPS antigens were modified by 4-(4,6-dimethoxy[1,3,5]triazin-2-yl)-4-methyl-morpholinium and conjugated to Luminex Magplex carboxylated beads. Antigen-coupled microspheres were washed and incubated with plasma samples at an appropriate sample dilution (1:1000 for all Isotypes and Fc-receptors) for 2 hours at 37°C in 384-well plates (Greiner Bio-One). Unbound antibodies were washed away, and antigen-bound antibodies were detected by using a PE-coupled detection antibody for each subclass and isotype (IgG1, IgG2, IgG3, IgA1, IgA2 and IgM; Southern Biotech), and Fc-receptors were fluorescently labeled with PE before addition to immune complexes (FcγR2A, FcγR2B, FcγR3A, FcγR3B, FcαR; Duke Protein Production facility). After one hour incubation, plates were washed, and flow cytometry was performed with an IQue (Intellicyt), and analysis was performed on IntelliCyt ForeCyt (v8.1). PE median fluorescent intensity (MFI) was reported as a readout for antigen-specific antibody titers.

#### Antibody-dependent monocyte and neutrophil phagocytosis (ADMP and ADNP)

ADMP and ADNP were conducted as previously described^9,43^. IpaB and IpaC were biotinylated using Sulfo-NHS-LC-LC biotin (Thermo Fisher) and coupled to fluorescent Neutravidin-conjugated beads (Thermo Fisher). *S. flexneri* 2a and *S. sonnei* OSP were modified with DMTMM and coupled to carboxylated fluorescent beads (Thermo Fisher). To form immune complexes antigen-coupled beads were incubated for 2 hours at 37°C with diluted samples (1:50) and then washed to remove unbound immunoglobulins. For ADMP, the immune complexes were incubated for 4 hours with fresh blood monocytes isolated with commercially available kit (StemCell) (1.25x10^5^ monocytes/mL) and for ADNP for 1 hour with fresh blood neutrophils isolated from healthy donors with commercially available kit (StemCell). For ADNP, neutrophils were washed and then fixed in 4% PFA. For ADMP, monocytes were washed, stained for CD14 (Biolegend), and in IgG depletion experiment for CD16, CD86, CX3CR1, CCR2 and HLA-DR and then fixed in 4% PFA. Flow cytometry was performed to identify the percentage of cells that had phagocytosed beads as well as the number of beads that had been phagocytosed (phagocytosis score = % positive cells × Median Fluorescent Intensity of positive cells/10000). Flow cytometry was performed with an IQue (Intellicyt) or BD LSR Fortessa, and analysis was performed on IntelliCyt ForeCyt (v8.1) or using FlowJo V10.7.1.

#### Antibody dependent complement deposition (ADCD)

ADCD was conducted using a 384-well based customized multiplexed assay. Protein antigens were coupled to magnetic Luminex beads (Luminex Corp) by carbodiimide-NHS ester-coupling (Thermo Fisher). OSP and LPS antigens were modified by 4-(4,6-dimethoxy[1,3,5]triazin-2-yl)-4-methyl-morpholinium and conjugated to Luminex Magplex carboxylated beads^44^. To form immune complexes, antigen-coupled beads were incubated for 2 hours at 37°C with diluted samples (1:50) and then washed to remove unbound immunoglobulins. Lyophilized guinea pig complement (Cedarlane) was resuspended according to manufacturer’s instructions and diluted in gelatin veronal buffer with calcium and magnesium (Boston BioProducts). Resuspended guinea pig complement was added to immune complexes and incubated for 20 minutes at 37°C. Post incubation, C3 was detected with Fluorescein-Conjugated Goat IgG Fraction to Guinea Pig Complement C3 (Mpbio).

#### IgA and IgG depletion

IgA or IgG were depleted from human plasma samples using CaptureSelect™ IgA or IgG Affinity Matrix (Thermo Fisher). The capture matrix was washed three times with PBS and incubated over night with 1:5 diluted plasma samples in a low protein binding MultiScreen® filter plate (Millipore). Depleted plasma was recovered by centrifugation of the filter plate. For each sample, non-depleted plasma was treated similarly but without affinity matrix.

#### Data Pre-processing

The raw MFI was scaled by the log10 function and was then subtracted by the corresponding PBS values. The normalized MFI values were assigned to zero if they were negative.

#### Computational analysis

A supervised multivariate analysis method of Least Absolute Shrinkage and Selection Operator (LASSO)^45^ followed by Partial Least Squares Discriminant Analysis(PLS-DA) was used to identify key antibody features that contribute to variation in the disease severity. Prior to building the LASSO-PLSR model, all titer, FcR and ADCD measurements were log transformed, and all measurements were then z-scored. LASSO identified a minimal set of features that drives separation in samples of endemic or non-endemic exposure and varying disease severity. To estimate the minimal correlates that best explained group differences without overfitting, a five-fold nested validation framework was designed. In each repetition, the dataset was randomly divided into groups of 5 randomly assorted individuals, where 80% of the dataset was used for building the model and the remaining holdout set was used to test the model prediction, where the goodness-of-fitness of the model was measured by classification accuracy between groups. At each LASSO run lambda parameter was optimized using *cv.glmnet* function. Ultimately, this approach resulted in the generation of a model with the minimal set of features that generated the best classification prediction in a cross-validation test. LASSO selected features were used to build the PLS-DA model regressing against the disease severity score. The performance of the algorithm was evaluated with R^2^ and Q^2^ metrics. Features were ranked based on their Variable of Importance (VIP) score and the loadings of the latent variable 1 (LV1) was visualized in a bar graph, which captures the contribution of each feature to the variation in disease severity. These analyses were carried out using R package *glmnet* (v4.0.2)^46^ and *ropls* (v1.20.0)^47^. For both *S.flexneri* 2a and *S.sonnei* infected over and under age 5 PLSDA model accuracy was 0.78. Co-correlate networks were constructed based on the pairwise correlation between the top predictive features selected and all measured biophysical and functional features. Only correlations with an absolute Spearman correlation coefficient greater than 0.7 and *p*-value lower than 0.01 after correction for multiple comparisons by Benjamini-Hochberg (BH) were shown. Networks were generated using R package *network* (v1.16.0) [Butts C (2015). network: Classes for Relational Data.].

### Statistical analysis

Microsoft Excel was used to compile and annotate experimental data. All plots were generated in Graph Pad Prism V.8. Statistical differences between two groups were calculated using a two-sided Mann–Whitney U test or Wilcoxon signed-rank test. To compare multiple groups, a Kruskal–Wallis test or Friedman test was used followed by the Dunn’s method correcting for multiple comparisons in Graph Pad Prism. Heat maps and correlation matrix were created using R version 1.4.1106. Spearmen rank correlation coefficient was used in all analysis.

## Supporting information

SuppTable1

## Acknowledgments

This study was approved by the Institutional Review Board of the International Centre for Diarrhoeal Disease Research, Bangladesh’s and from the Massachusetts General Hospital’s Institutional Review Board. We obtained written documentation of informed consent from all adult participants and from guardians of participating children before enrollment.

This research was supported by the icddr,b and extramural grants from the National Institutes of Health, including the National Institute of Allergy and Infectious Diseases (NIAID) (R01AI155414, R01AI177075) and the Fogarty International Center and NIAID Training Grant in Vaccine Development and Public Health (D43 TW005572; SRB, MAA, SM, NHR). The International Centre for Diarrhoeal Disease Research, Bangladesh (icddr,b) is thankful to the donors for their support of its research efforts. icddr,b is also grateful to the governments of Bangladesh and Canada for providing core/unrestricted support. SC was also supported by National Institute of Allergy and Infectious Diseases of the NIH under Award No. R01AI153399. BB is the recipient of an HSFP postdoctoral fellowship, a Zuckerman STEM Leadership Program fellowship and the Weizmann Institute of Science Women’s Postdoctoral Career Development Award.

## Supp Figures

**Supp Fig 1.**
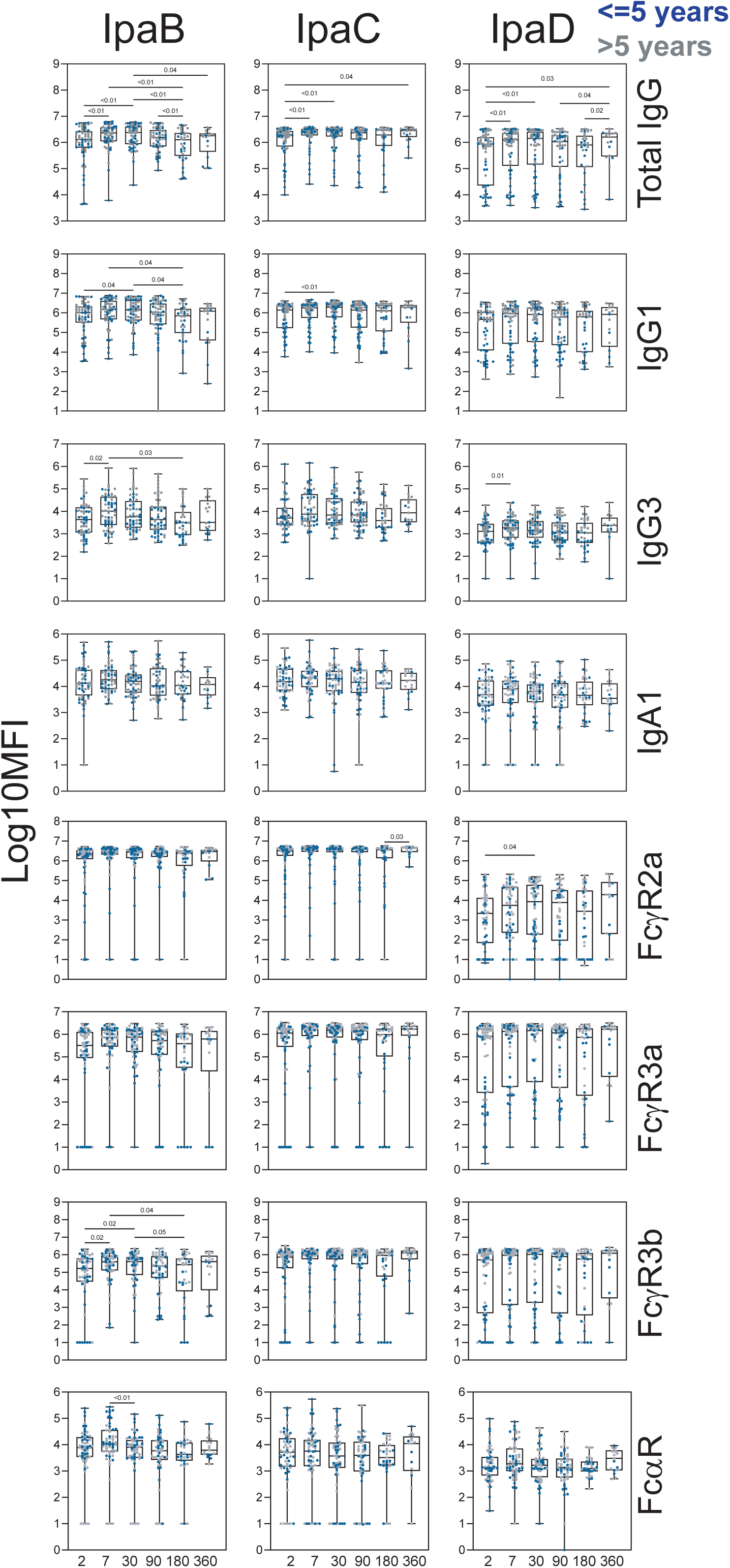
Shigella protein-specific antibody responses to shigellosis. *Shigella* protein-specific total IgG, IgG1, IgG3, IgA1 and binding to FcγR2a, FcγR3a, FcγR3b and FcαR in serum of individuals presenting with stool culture-confirmed shigellosis in Bangladesh. Horizontal line depicts median. (p-value calculated by Mixed effects analysis followed by Holm-Sidak’s multiple comparisons test)

**Supp Fig 2.**
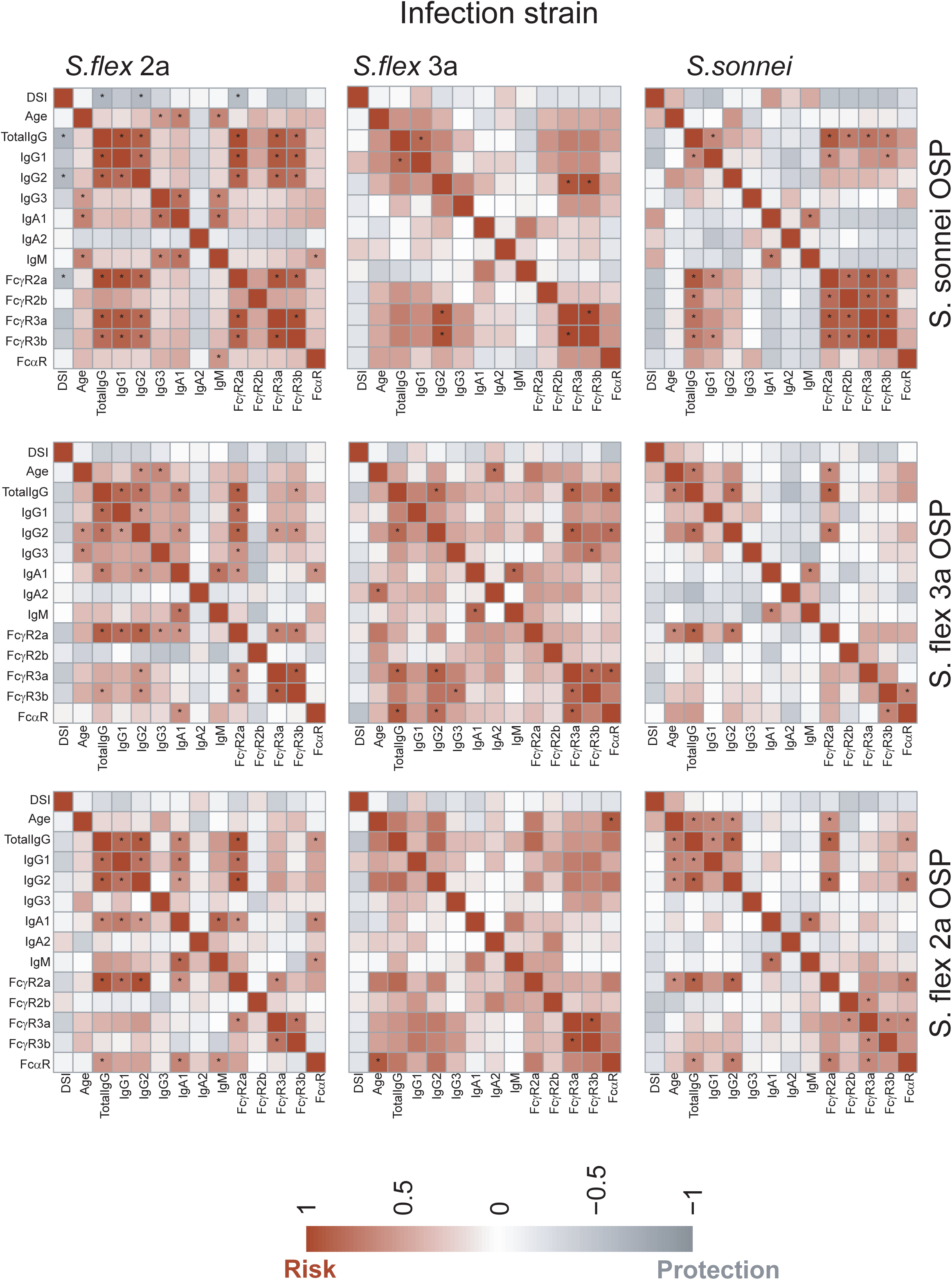
OSP-specific antibody-mediated cross protection against shigellosis. Heat map of Spearman correlation of disease severity and age with Shigella OSP-specific antibody features in serum of individuals presenting with *S. flexneri* 2a, *S. flexneri* 3a or *S. sonnei* stool culture-confirmed shigellosis on day 2 of clinical care. (*Benjamini-Hochberg corrected p-value<0.05)

**Supp Fig 3.**
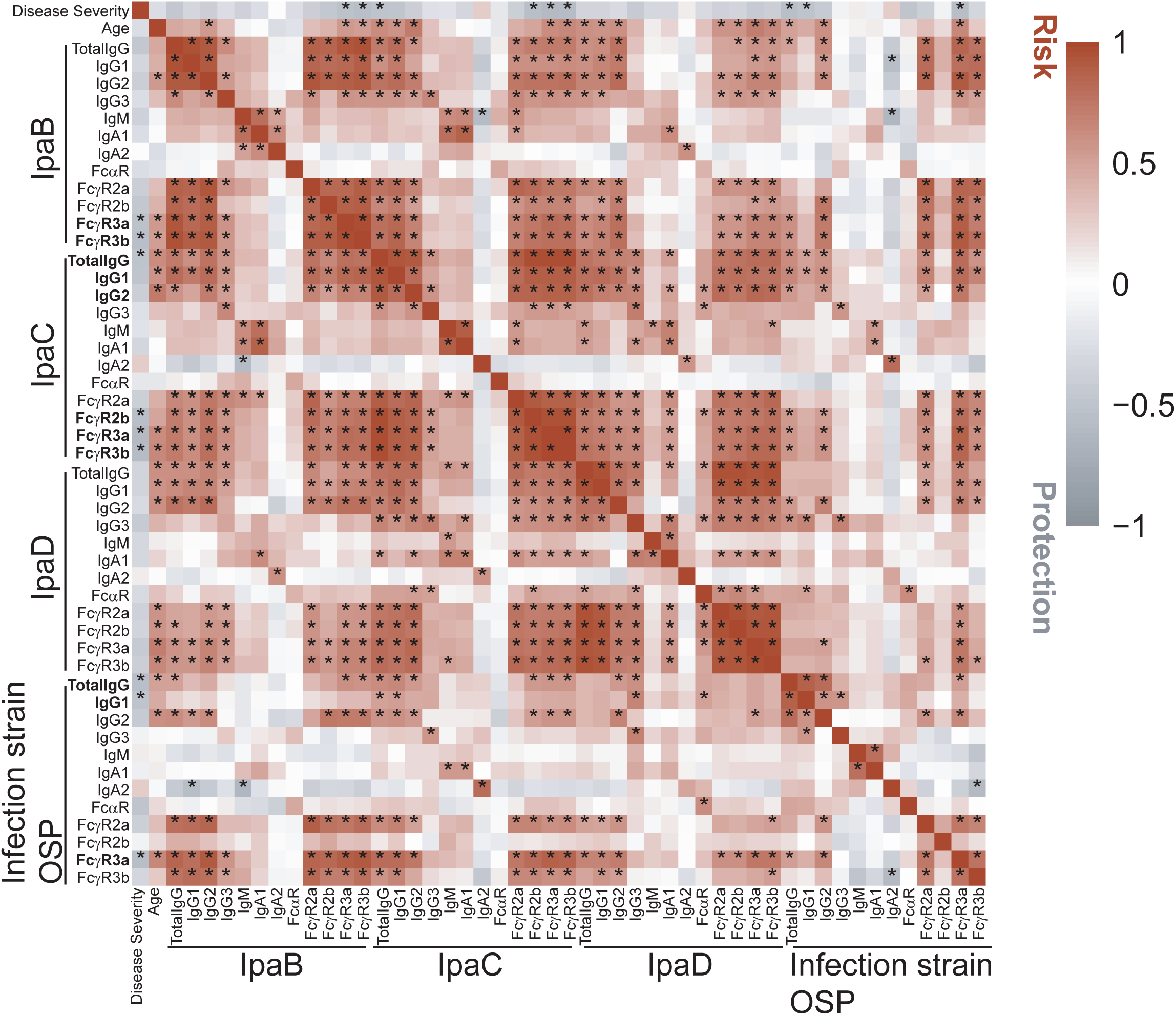
Impact of antibody profiles on disease severity for a subset of patients presenting for clinical care up to 48hrs after symptom onset. Heat map of Spearman correlation of disease severity and age with Shigella-specific antibody features in serum of individuals presenting with stool culture-confirmed shigellosis on day 2 of clinical care, up to 48hrs since symptom onset. (*Benjamini-Hochberg corrected p-value<0.05)

**Supp Fig 4.**
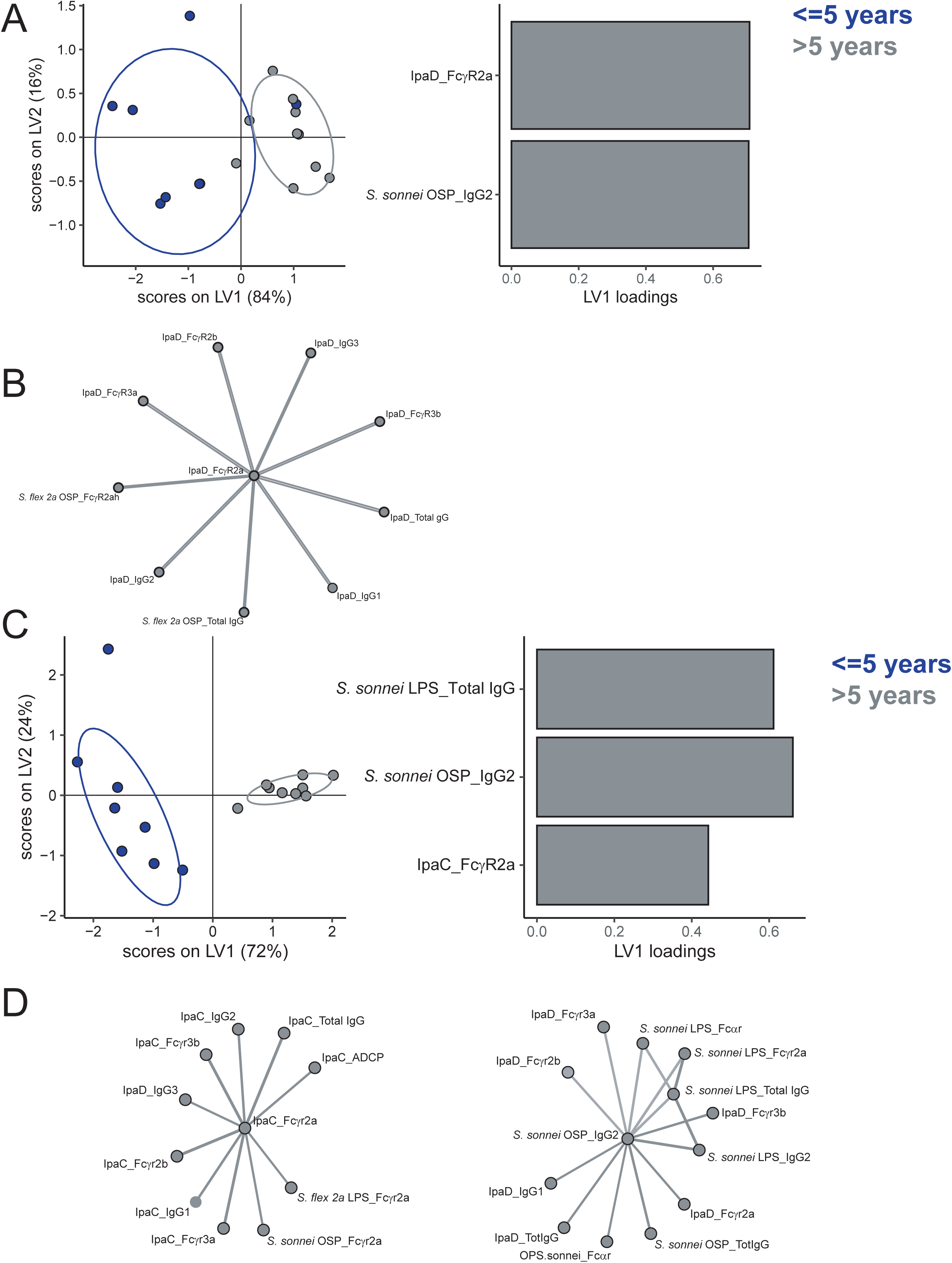
Age-dependent differences in immune response to shigellosis. Score plots and LV1 loadings of PLS-DA model using LASSO selected features of Shigella-specific antibody isotypes and FcR binding in serum of individuals on day 7 (A-B) or day 90 (C-D) after clinical presentation with stool culture-confirmed shigellosis caused by *S. sonnei* and corresponding correlation networks of selected antibody features(B,D) (Spearman rank coefficient R>0.7, Benjamini-Hochberg corrected p<0.01)

## References

1. Black, R. E. et al. Estimated global and regional causes of deaths from diarrhoea in children younger than 5 years during 2000–21: a systematic review and Bayesian multinomial analysis. Lancet Glob Health 12, e919–e928 (2024).

2. Kotloff, K. L. et al. Burden and aetiology of diarrhoeal disease in infants and young children in developing countries (the Global Enteric Multicenter Study, GEMS): a prospective, case-control study. 382, 209–222 (2013).

3. Cohen, D. et al. Serum IgG antibodies to Shigella lipopolysaccharide antigens–a correlate of protection against shigellosis. Hum Vaccin Immunother 15, 1401–1408 (2019).

4. Simon, A. K., Hollander, G. A. & McMichael, A. Evolution of the immune system in humans from infancy to old age. Proceedings of the Royal Society B: Biological Sciences 282, (2015).

5. Shimanovich, A. A. et al. Functional and antigen-specific serum antibody levels as correlates of protection against shigellosis in a controlled human challenge study. Clinical and Vaccine Immunology 24, (2017).

6. Bernshtein, B. et al. Systems approach to define humoral correlates of immunity to Shigella. Cell Rep 40, 111216 (2022).

7. Cohen, D., Green, M. S., Block, C., Slepon, R. & Ofek, I. Prospective study of the association between serum antibodies to lipopolysaccharide O antigen and the attack rate of shigellosis. J Clin Microbiol 29, 386–389 (1991).

8. Cohen, D., Block, C., Green, M. S., Lowell, G. & Ofek, I. Immunoglobulin M, A, and G antibody response to lipopolysaccharide O antigen in symptomatic and asymptomatic Shigella infections. J Clin Microbiol 27, 162–167 (1989).

9. Bernshtein, B. et al. Determinants of immune responses predictive of protection against shigellosis in an endemic zone: a systems analysis of antibody profiles and function. Lancet Microbe 100889 (2024) doi:10.1016/S2666-5247(24)00112-5.

10. MacLennan, C. A., Grow, S., Ma, L. F. & Steele, A. D. The Shigella Vaccines Pipeline. Vaccines (Basel) 10, (2022).

11. Lu, T., Das, S., Howlader, D. R., Picking, W. D. & Picking, W. L. Shigella Vaccines: The Continuing Unmet Challenge. Int J Mol Sci 25, (2024).

12. Giersing, B. K. et al. Clinical and regulatory development strategies for Shigella vaccines intended for children younger than 5 years in low-income and middle-income countries. Lancet Glob Health 11, e1819–e1826 (2023).

13. Passwell, J. H. et al. Age-related efficacy of Shigella O-specific polysaccharide conjugates in 1-4-year-old Israeli children. Vaccine 28, 2231–2235 (2010).

14. Porter, C. K. et al. Clinical endpoints in the controlled human challenge model for Shigella: A call for standardization and the development of a disease severity score. PLoS One 13, (2018).

15. Robin, G. et al. Characterization and quantitative analysis of serum IgG class and subclass response to Shigella sonnei and Shigella flexneri 2a lipopolysaccharide following natural Shigella infection. Journal of Infectious Diseases 175, 1128–1133 (1997).

16. Cohen, D., Green, M. S., Biock, C., Rouach, T. & Ofek, I. Serum antibodies to lipopolysaccharide and natural immunity to shigellosis in an israeli military population. Journal of Infectious Diseases 157, 1068–1071 (1988).

17. Martinez-Becerra, F. J. et al. Broadly protective Shigella vaccine based on type III secretion apparatus proteins. Infect Immun 80, 1222–1231 (2012).

18. Kotloff, K. L. The Burden and Etiology of Diarrheal Illness in Developing Countries. Pediatr Clin North Am 64, 799–814 (2017).

19. Kotloff, K. L., et al. Global Burden of Shigella Infections: Implications for Vaccine Development and Implementation of Control Strategies. Bulletin of the World Health Organization vol. 77 651–666 (World Health Organization, 1999).

20. Kotloff, K. L. et al. Burden and aetiology of diarrhoeal disease in infants and young children in developing countries (the Global Enteric Multicenter Study, GEMS): a prospective, case-control study. 382, 209–222 (2013).

21. Lowell, G. H. et al. Antibody-dependent cell-mediated antibacterial activity: K lymphocytes, monocytes, and granulocytes are effective against shigella. The Journal of Immunology 125, 2778–2784 (1980).

22. Hathaway, L. J., Griffin, G. E., Sansonetti, P. J. & Edgeworth, J. D. Human monocytes kill Shigella flexneri but then die by apoptosis associated with suppression of proinflammatory cytokine production. Infect Immun 70, 3833–3842 (2002).

23. Schnupf, P. & Sansonetti, P. J. Shigella Pathogenesis: New Insights through Advanced Methodologies. Microbiol Spectr 7, (2019).

24. Zohar, T. et al. A multifaceted high-throughput assay for probing antigen-specific antibody-mediated primary monocyte phagocytosis and downstream functions. J Immunol Methods 510, 113328 (2022).

25. Cohen, D. et al. Safety and immunogenicity of investigational Shigella conjugate vaccines in Israeli volunteers. Infect Immun 64, 4074–4077 (1996).

26. Ashkenazi, S. & Cohen, D. An update on vaccines against Shigella . Ther Adv Vaccines 1, 113–123 (2013).

27. Levine, M. M., Kotloff, K. L., Barry, E. M., Pasetti, M. F. & Sztein, M. B. Clinical trials of Shigella vaccines: Two steps forward and one step back on a long, hard road. Nat Rev Microbiol 5, 540– 553 (2007).

28. Fernandez-Prada, C. M. et al. Shigella flexneri IpaH(7.8) facilitates escape of virulent bacteria from the endocytic vacuoles of mouse and human macrophages. Infect Immun 68, 3608–3619 (2000).

29. Way, S. S., Borczuk, A. C., Dominitz, R. & Goldberg, M. B. An Essential Role for Gamma Interferon in Innate Resistance to Shigella flexneri Infection. Infect Immun 66, 1342 (1998).

30. Lemme-Dumit, J. M., Doucet, M., Zachos, N. C. & Pasetti, M. F. Epithelial and Neutrophil Interactions and Coordinated Response to Shigella in a Human Intestinal Enteroid-Neutrophil Coculture Model. mBio 13, (2022).

31. François, M., Le Cabec, V., Dupont, M. A., Sansonetti, P. J. & Maridonneau-Parini, I. Induction of necrosis in human neutrophils by Shigella flexneri requires type III secretion, IpaB and IpaC invasins, and actin polymerization. Infect Immun 68, 1289–1296 (2000).

32. Mandic-Mulec, I., Weiss, J. & Zychlinsky, A. Shigella flexneri is trapped in polymorphonuclear leukocyte vacuoles and efficiently killed. Infect Immun 65, 110–115 (1997).

33. Reed, W. P. Serum factors capable of opsonizing Shigella for phagocytosis by polymorphonuclear neutrophils. Immunology 28, 1051 (1975).

34. Chakraborty, S., Connor, S. & Velagic, M. Development of a simple, rapid, and sensitive diagnostic assay for enterotoxigenic E. coli and Shigella spp applicable to endemic countries. PLoS Negl Trop Dis 16, (2022).

35. Chowdhury, G. et al. Field evaluation of a simple and rapid diagnostic test, RLDT to detect Shigella and enterotoxigenic E. coli in Indian children. Sci Rep 14, (2024).

36. Brown, E. P. et al. Optimization and qualification of an Fc Array assay for assessments of antibodies against HIV-1/SIV. J Immunol Methods 455, 24–33 (2018).

37. Kelly, M. et al. Development of Shigella conjugate vaccines targeting Shigella flexneri 2a and S. flexneri 3a using a simple platform-approach conjugation by squaric acid chemistry. Vaccine 41, 4967–4977 (2023).

38. Xu, P. et al. Simple, direct conjugation of bacterial O-SP-core antigens to proteins: development of cholera conjugate vaccines. Bioconjug Chem 22, 2179–2185 (2011).

39. Kováč, P. & Xu, P. Controlled and highly efficient preparation of carbohydrate-based vaccines: Squaric acid chemistry is the way to go. Carbohydr Chem 42, 83–115 (2017).

40. WO 2013/009826 A1 - Conjugating Amines | The Lens. https://www.lens.org/lens/patent/044-485-774-710-28X/frontpage?l=en.

41. Xu, P. et al. Conjugate Vaccines from Bacterial Antigens by Squaric Acid Chemistry: A Closer Look. Chembiochem 18, 799–815 (2017).

42. Kelly, M. et al. Development of a Shigella conjugate vaccine targeting Shigella flexneri 6 that is immunogenic and provides protection against virulent challenge. Vaccine 42, 126263 (2024).

43. Butler, A. L., Fallon, J. K. & Alter, G. A sample-sparing multiplexed ADCP assay. Front Immunol 10, 1851 (2019).

44. Schlottmann, S. A. et al. A novel chemistry for conjugating pneumococcal polysaccharides to Luminex microspheres. 309, 75–85 (2006).

45. Tibshirani, R. The lasso method for variable selection in the cox model. Stat Med 16, 385–395 (1997).

46. Friedman, J., Hastie, T. & Tibshirani, R. Regularization paths for generalized linear models via coordinate descent. J Stat Softw 33, 1–22 (2010).

47. Thévenot, E. A., Roux, A., Xu, Y., Ezan, E. & Junot, C. Analysis of the Human Adult Urinary Metabolome Variations with Age, Body Mass Index, and Gender by Implementing a Comprehensive Workflow for Univariate and OPLS Statistical Analyses. J Proteome Res 14, 3322– 3335 (2015).

