## Supplementary material for "Limited O-specific polysaccharide (OSP)-specific functional antibody responses in young children with Shigella infection in Bangladesh": SuppTable1

| **Infection Strain** | **Total # of patients** | **# of patients <= age 5** | **# of patients**  **> age 5** | **% Males** | **%Blood in diarrhea** | **%Fever** | **%Vomiting** | **Average number of stools (SD)** | **Average duration of diarrhea in hours (SD)** | **Average DSI (SD)** |
| --- | --- | --- | --- | --- | --- | --- | --- | --- | --- | --- |
| *S. flexneri* 2a | 20 | 8 | 12 | 50 | 95 | 50 | 25 | 6.5 (3.6) | 72.7 (61.6) | 3.7 (1.2) |
| *S. flexneri* 3a | 11 | 7 | 4 | 63.6 | 100.0 | 72.7 | 45.5 | 5.2 (3.5) | 79.8 (29.5) | 4 (0.7) |
| *S. flexneri* 6 | 5 | 2 | 3 | 60.0 | 100.0 | 0.0 | 0.0 | 2.3 (0.4) | 95.2 (38.7) | 1.8 (0.4) |
| *S. Sonnei* | 19 | 8 | 11 | 47.4 | 100.0 | 52.6 | 10.5 | 8.4 (4.0) | 69.6 (51.1) | 3.7 (1.1) |

**Supp Table 1. Study design and participants.**
